# AP-1 mediated chromatin changes govern alveolar type 2 cell transition in lung injury-repair

**DOI:** 10.1101/2025.10.25.684549

**Authors:** Anne M Lynch, Tala Noun, Sujuan Yang, Tieling Zhou, Scott E Evans, Humam Kadara, Jichao Chen

**Affiliations:** Department of Pulmonary Medicine, the University of Texas MD Anderson Cancer Center, Houston, Texas 77030, USA; Graduate Program in Developmental Biology, Baylor College of Medicine, Houston, TX, USA; Department of Translational Molecular Pathology, The University of Texas MD Anderson Cancer Center, Houston, TX, USA; The University of Texas MD Anderson Cancer Center UTHealth Houston Graduate School of Biomedical Sciences, Houston, TX, USA; Department of Pediatrics, Perinatal Institute Division of Pulmonary Biology, University of Cincinnati and Cincinnati Children’s Hospital Medical Center, Cincinnati, OH 45229, USA

**Keywords:** Lung injury-repair, transitional AT2 cells, AP-1, epigenomics

## Abstract

Facultative stem cells including lung alveolar type 2 (AT2) cells must toggle between unipotent specialization at baseline and multipotent plasticity during injury-repair. The underlying molecular switches and epigenetic changes remain unclear yet dictate regenerative versus pathological outcomes. Using multiomics and mouse genetics, we show that AP-1 members FOS, FOSB, and JUNB promote an injury-induced transitional AT2 cell state and associated chromatin landscape. Joint transcriptomic-epigenomic profiling and immunostaining distinguish CLDN4+ AT2 cells as a KRT8^high^ subset with open chromatin highly enriched for AP-1 motifs. JUNB and FOSB accumulate in these cells upon viral injury and, along with constitutive FOS, are required for CLDN4 induction, AT2 cell dispersion, and senescence signaling toward fibroblasts, while impacting region-specific alveolar type 1 (AT1) differentiation. AP-1 activation also occurs in mouse AT2 cells expressing oncogenic Kras and transitional cells in human lung tissues with premalignant or adenocarcinoma lesions. Our work refines AT2 transitional states and reveals a gene regulatory logic shared by tissue repair and tumorigenesis.

**SIGNIFICANCE STATEMENT:** Facultative stem cells balance between their resting mode and regenerative mode. In the lung, alveolar type 2 (AT2) cells produce surfactants at baseline and, upon injury, activate to self-renew and differentiate into alveolar type 1 (AT1) cells. Such activation, observed in acute lung injury, fibrosis, and tumorigenesis, proceeds through a high-KRT8 transitional phase. Using multiomic profiling and mouse genetics, we identify activator protein-1 (AP-1) transcription factors as central regulators of a CLDN4⁺ transitional substate, promoting AT2 cell dispersion, fibroblast signaling, and region-specific AT1 differentiation. AP-1 activation is conserved during oncogenic transformation in both mouse and human lungs. These findings refine our understanding of AT2 cell fate transition and chromatin dynamics, and reinforce a molecular connection between alveolar repair and tumorigenesis.

## INTRODUCTION

Resting facultative stem cells are unipotent and highly specialized—for example, surfactant-producing alveolar type 2 (AT2) cells in the lung, exocrine acinar cells in the pancreas, and bile-secreting cholangiocytes in the liver. Activated during homeostatic turnover or after injury, these cells self-renew and differentiate into AT1 cells, ductal cells, and hepatocytes, respectively—thus fulfilling the operational definition of stem cells ^1–4^. In the resting mode, cell-type and lineage-defining gene networks enforce their specialized functions. Triggered by physicochemical niche cues, the stem-cell mode is more dynamic and inherently unstable because concurrent processes of inflammation, barrier restoration, proliferation, migration, and differentiation tug in competing directions and fluctuate over time, while resisting and returning to the resting mode as needed. Orderly entry into and exit from the activated stem-cell mode requires choreographed gene regulation, which often falters in fibrotic and neoplastic diseases ^5–7^.

Resting AT2 cells are maintained by transcription factors such as NKX2-1, CEBPA, and ETV5 ^8–11^. Upon activation, this circuit must attenuate and be progressively supplanted by a network that retains NKX2-1 but now enlists YAP/TAZ/TEAD to drive AT1 differentiation ^8^. Complicating this transition, AT2 cells traverse a transient KRT8^high^ intermediate— documented during development, repair, and pathology ^12–16^ —that resembles the high-energy transition state of a chemical reaction. This intermediate, variously termed damage-associated transient progenitors (DATPs), pre-alveolar type-1 transitional cell state (PATS), or alveolar differentiation intermediate (ADI), exhibits overlapping yet distinct transcriptional signatures, underscoring functional heterogeneity needed to coordinate the said concurrent reparative programs ^12,15,17^. This transitional population is uniquely vulnerable, acquiring a pro-fibrotic, senescence-associated secretory phenotype (SASP) and seeding adenocarcinomas ^18,19^. Such biological complexity and medical relevance have spurred extensive research implicating mechanical (YAP/TAZ, CDC42), morphogenic (WNT, BMP, NOTCH), and immunological (IL1B) cues in governing AT2, AT1, and the intervening states, as well as the switches between them ^17,20–22^. Nevertheless, finer resolution of transitional-state heterogeneity remains essential to pinpoint the specific targets and downstream consequences of these and other regulators.

Moving beyond the transcriptional landscape to identify regulators, we employ bulk ATAC-seq and single-cell multiome profiling to resolve multiple AT2 transitional states after Sendai-virus injury. AP-1 is the most enriched motif during AT2 activation and reaches its highest accessibility in the CLDN4+ transitional subset. Deletion of *Fos*, *Fosb*, and *Junb* in AT2 cells closes transitional-state chromatin, blocks CLDN4 expression, limits AT2 dispersion and SASP-driven fibroblast activation, and impacts AT1 differentiation in a region-specific manner. Moreover, active AP-1 marks transitional AT2 cells in a Kras-driven lung-cancer mouse model and in human lung premalignant and adenocarcinoma lesions.

## RESULTS

### AP-1 motifs dominate reversible chromatin opening upon Sendai virus infection

To extend published transcriptomic analyses ^12,15,17,23–26^ and pinpoint action sites of transcriptional regulators, we performed bulk ATAC-seq on *Sftpc^CreER^* lineage–traced AT2 cells at baseline and at 14- and 49-day post–Sendai virus infection (dpi)—corresponding to peak proliferation and their subsequent return to quiescence, respectively ^27^ (Fig. 1A). Peaks with increased accessibility at 14 dpi were highly enriched for AP-1 motifs alongside ELF and ETV motifs, whereas those losing accessibility at 14 dpi were enriched for NKX and FOXA motifs, possibly reflecting downregulation of the NKX2-1/FOXA1/FOXA2–driven surfactant program of the resting mode ^28–30^ (Fig. 1B; Table S1; Table S2). Comparison of 49 dpi versus 14 dpi revealed that decreased peaks remained AP-1–enriched, while increased peaks featured NKX, FOXA, and CTCF motifs, suggesting chromatin restoration and reorganization (Fig. 1B). To follow individual peaks over time, we plotted fold changes for 14 dpi vs. baseline against 49 dpi vs. 14 dpi, showing remarkable clustering of AP-1– containing peaks in the quadrant indicative of transient opening at 14 dpi followed by closure at 49 dpi (Fig. 1C; Table S3).

**Fig. 1.**
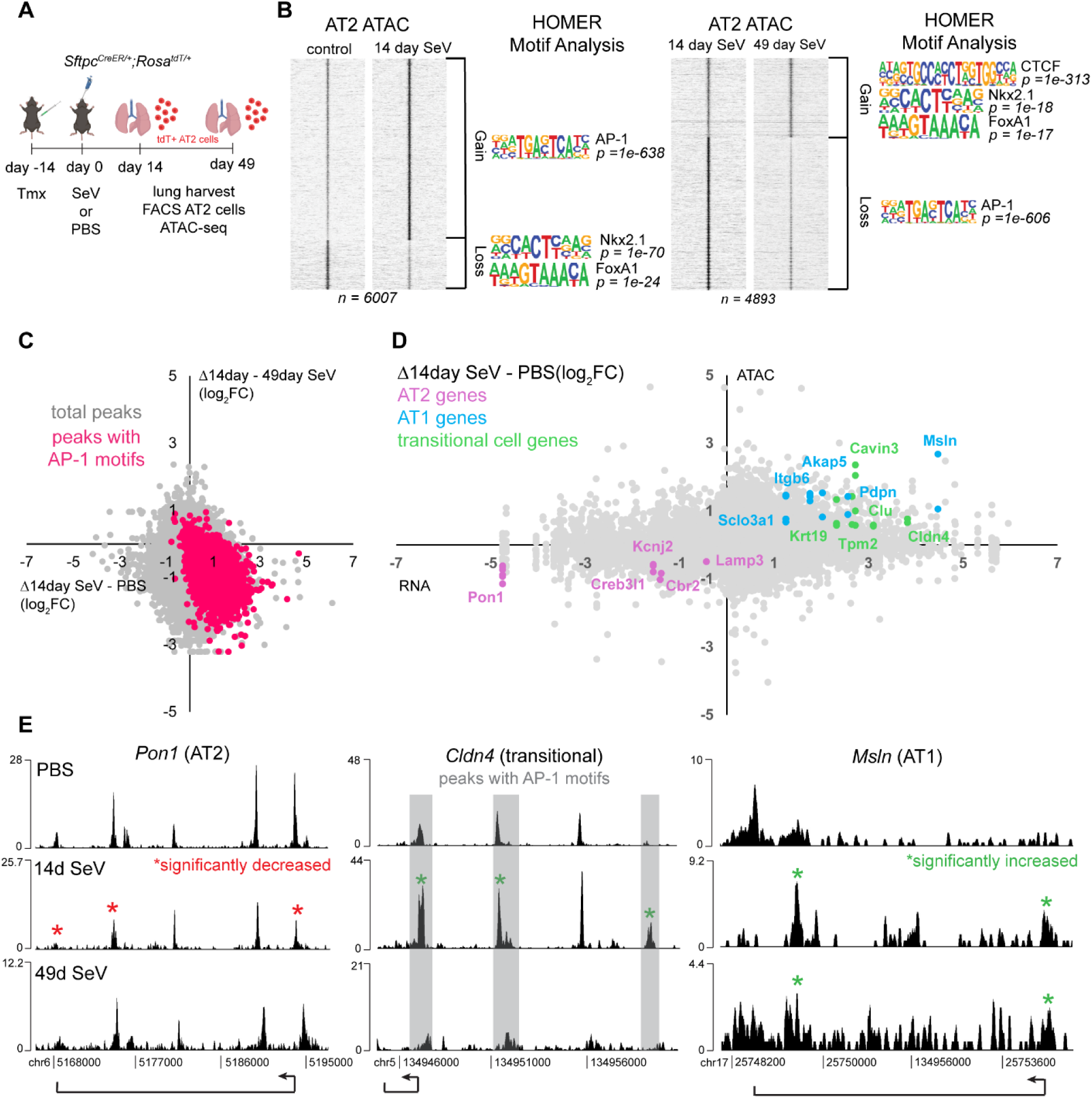
AP-1 motifs dominate reversible chromatin opening upon Sendai virus (SeV) infection. (**A**) Schematic of AT2 cell lineage trace, infection, and lung harvest for bulk ATAC-seq. (**B**) Heatmap of bulk ATAC-seq gained peaks (Log_2_FC > 1, pval < 0.05) and lost peaks (log_2_FC < −1, pval < 0.05) from 14-day SeV vs. PBS (left) and from 49-day SeV vs. 14-day SeV (right) with top HOMER motif analysis results displayed adjacent to each peak set. (**C**) Scatter plot of Log_2_FC changes of each ATAC-seq peak matched from 14-day SeV vs. PBS (x-axis) to 49-day SeV vs 14-day Sev (y-axis) to track the accessibility of the same peak regions over time. Peaks containing AP-1 motifs are highlighted in red and are mostly limited to the fourth quadrant. (**D**) Scatter plot of Log_2_FC of RNA expression (x-axis; pseudobulk single-cell RNA-seq) and ATAC accessibility (y-axis; bulk ATAC-seq peaks annotated to the nearest genes) of genes in AT2 cells comparing 14-day SeV and PBS lungs. Representative AT2 genes are highlighted in purple, AT1 genes in blue, and transitional cell genes in green. (**E**) Coverage plots of significantly changed peaks (asterisks) in genes representative of AT2 (*Pon1*), transitional (*Cldn4*), and AT1 (*Msln*) cell fate. Gray bars highlight peaks that contain one or more AP-1 motifs. Plot axes were normalized based on fraction of reads in each peak (FriP).

Linking peaks to their nearest genes revealed largely concordant changes upon viral infection in chromatin accessibility and previously reported RNA abundance from scRNA-seq ^26^, albeit with a larger dynamic range for RNA (Fig. 1D; Table S4). Transient accessibility gains at 14 dpi were accompanied by induction of transitional AT2 markers *Cldn4* and *Krt19* as well as AT1 genes *Pdpn* and *Msln*, alongside transient repression of core AT2 genes *Pon1* and *Lamp3* (Fig. 1D). By 49 dpi, AT2 genes such as *Pon1* regained accessibility and transitional markers such as *Cldn4* returned to baseline accessibility, whereas AT1 genes such as *Msln* maintained newly accessible peaks, reflecting AT1 differentiation of an AT2 subset (Fig. 1E). Thus, Sendai virus infection switches resting AT2 cells to the stem-cell mode, marked by reversible opening of AP-1 containing peaks and expression of associated genes.

### Single-cell multiome resolves four transitional AT2 populations including an AP-1 enriched CLDN4+ substate

To resolve AT2 chromatin dynamics at single-cell resolution and dissect the diverse AT2 activation programs, we performed single-cell multiome profiling on sorted CDH1+ epithelial cells from *Sftpc^CreER/+^; Rosa^Sun1GFP/+^* mice at 14 dpi with Sendai virus or PBS control. After excluding airway and Krt5+ dysplastic cells (Fig. S1), weighted joint analysis of RNA and ATAC profiles identified AT1 cells (*Rtkn2*+) and seven AT2 populations: canonical (*Lamp3*+), proliferating (*Mki67*+), chemokine-producing (*Ccl20*+), and four lineage-traced (Sun1GFP+) transitional substates (T1–T4) (Fig. 2A; Table S5). Module scoring revealed a progression from AT2 to AT1: T1, T3, and T4 showed stepwise gains in AT1 gene and chromatin signatures with concurrent loss of AT2 features (less dramatic for chromatin), whereas T2 exhibited minimal AT1 or AT2 scores but maximal enrichment for KRT8 ADI (alveolar differentiation intermediate) and PATS (pre-alveolar type-1 transitional cell state) scores, including specific *Cldn4* expression (Fig. 2B, 2A; Table S6) ^12,15^. The ADI score was low in AT1 cells, whereas the PATS score remained elevated, underscoring their emphasis on AT2 activation and AT1 differentiation, respectively (Table S5). Notably, although the AP-1 member RNA score was variable, AP-1 motif accessibility was elevated upon infection and spiked in T2, identifying the primary contributor to AP-1 motif enrichment in bulk ATAC (Fig. 2B, 2A). Congruently, each population had its characteristic motif enrichment with AP-1 motif marking T2 and AT1-specific TEAD motif marking T1, T3, and T4 (Table S7).

**Fig. 2.**
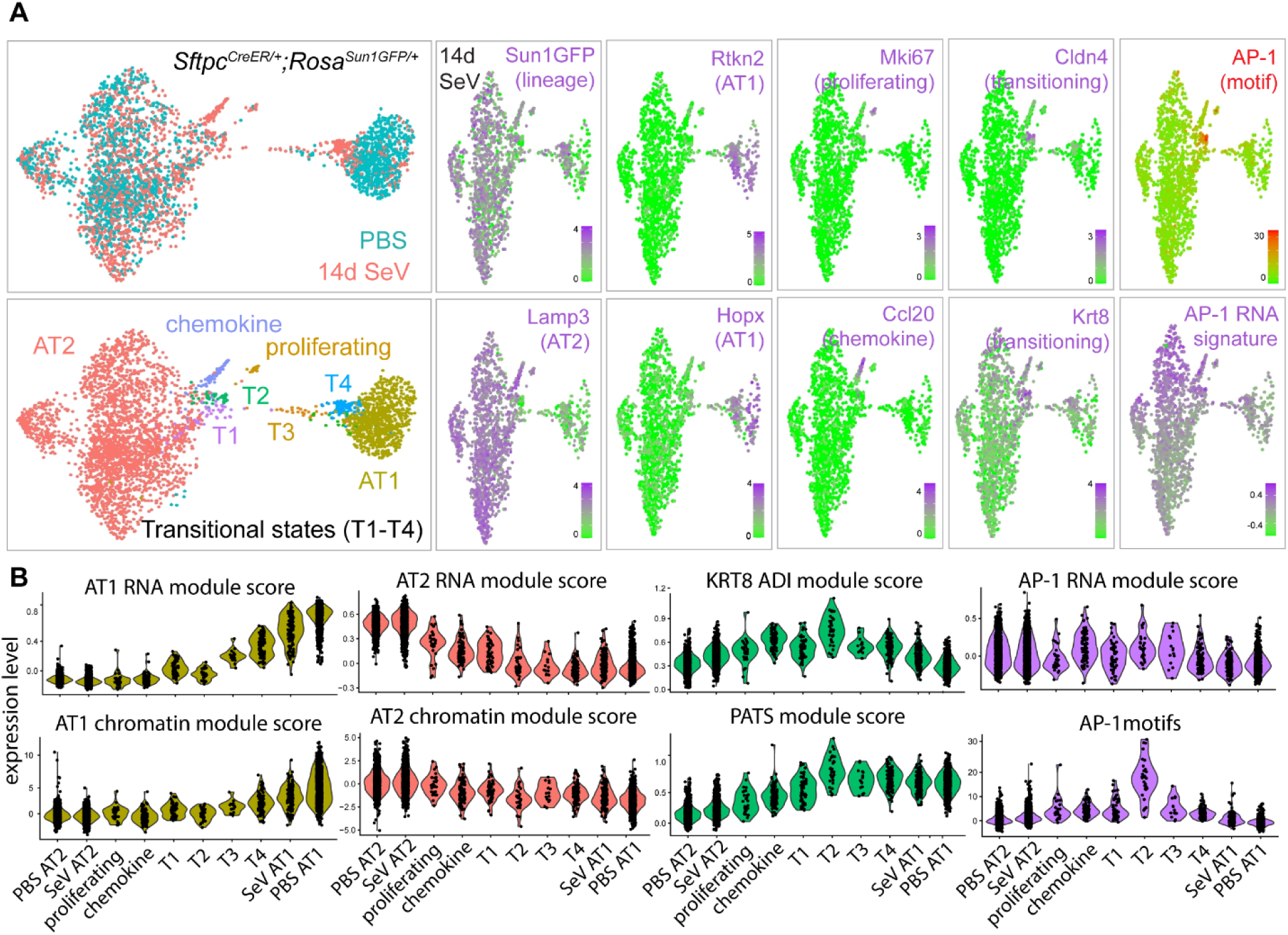
Single-cell multiome resolves four transitional AT2 populations including an AP-1 enriched CLDN4+ substate. (**A**) Single-cell multiome (ATAC+RNA) weighted nearest neighbor UMAPs of AT1 and AT2 cells, colored for PBS control and 14-day SeV infected lungs (top) and 7 AT2 cell populations (bottom). Feature plots of SeV infected lungs with representative gene expression from each cluster (purple), AP-1 gene module score (purple), and AP-1 motifs (red). (**B**) Violin plots across UMAP populations of RNA module scores using published gene sets specific to AT2 cells, AT1 cells, KRT8 ADI, PATS, and AP-1 transcription factors; AT2 and AT1 cell specific chromatin module scores generated from linked peaks of RNA modules; and a representative AP-1 motif.

To validate and spatially map the transitional substates identified by single-cell multiome, we selected informative, robust markers: LAMP3 for resting surfactant–producing AT2 cells; MHCII for AT2 identity; high KRT8 for activated AT2 cells (as well as normal airway cells); CLDN4 for the T2 substate; HOPX for differentiating AT2 cells (as well as AT1 cells); and *Rosa^tdT^* for lineage tracing and cell morphology. Single-cell quantitation and serial-section immunostaining showed that T1 upregulated KRT8 and downregulated LAMP3 without CLDN4 or HOPX expression; T3 retained high KRT8, lacked CLDN4 and LAMP3, and began expressing HOPX; and T4 exhibited baseline KRT8, robust HOPX, and flattened morphology while retaining MHCII—distinguishing them from lineage-traced AT1 cells; whereas T2 was uniquely CLDN4⁺KRT8^high^ with no LAMP3 or HOPX (Fig. 3A, 3B, 3C). These observations supported a model in which T1, T3, and T4 progress along an AT2-to-AT1 differentiation continuum, whereas T2 represents a distinct AP-1–marked branch (Fig. 3D).

**Fig. 3.**
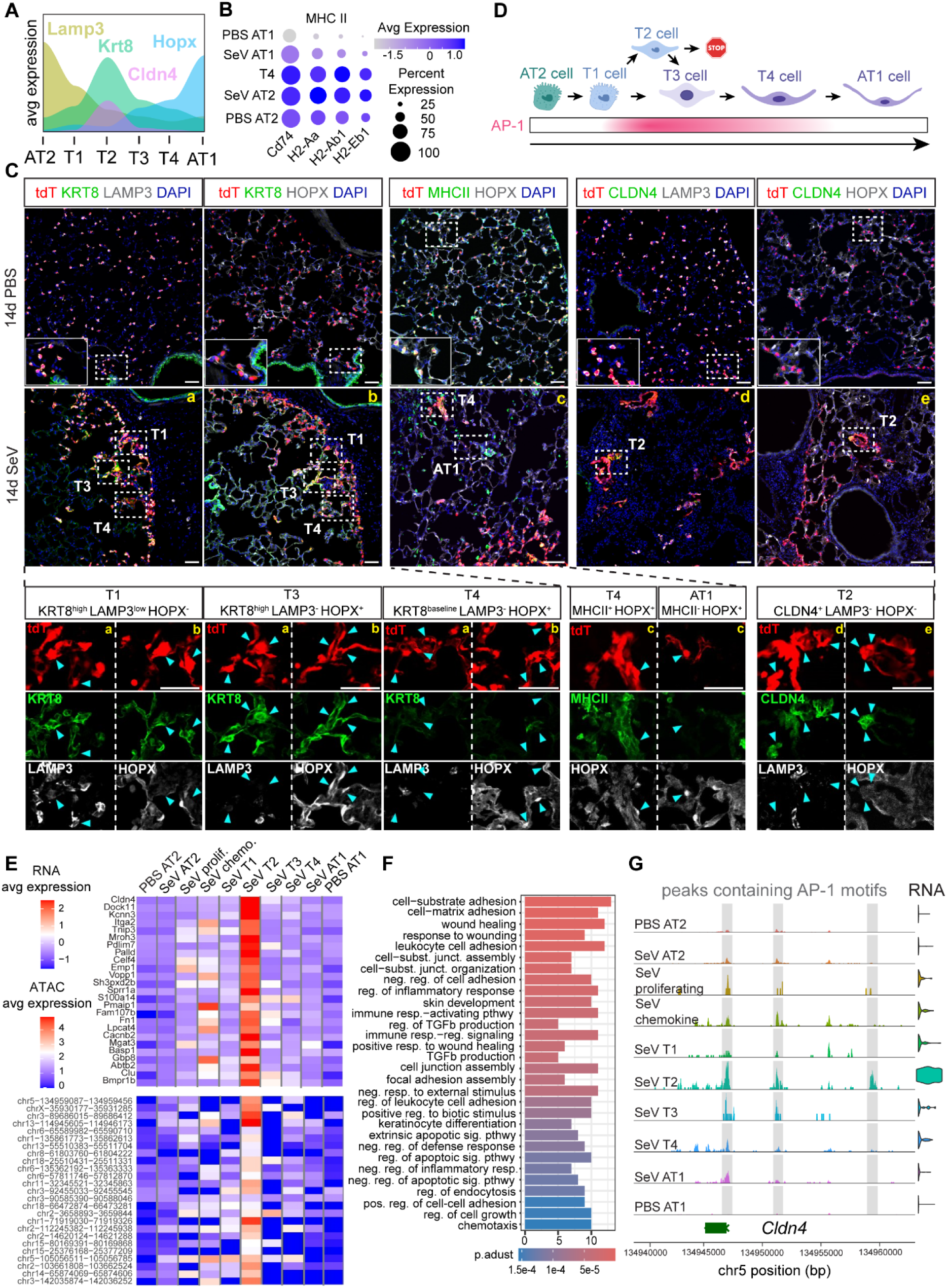
Immunostaining and molecular evidence for AT2 transitional substates. (**A**) Single-cell multiome average expression area plot of genes for AT2 cells (Lamp3), transitional cells (Krt8 and Cldn4), and AT1 cells (Hopx) within the transitional substates. (**B**) Single-cell multiome dotplot of genes that constitute MHCII molecules. Loss of MHCII distinguishes AT2 cells that are still in transition from those that have become AT1 cells. (**C**) Confocal images of 10µm sectioned lungs from PBS or SeV infected *Sftpc^CreER/+^;Rosa^tdT/+^*mice immunostained to identify the four transitional substates. Images a and b along with c and d are serial sections. Dashed line boxes are 2X magnification insets displayed on the bottom row and arrows point to cells indicative of each transitional substate (T1-T4). (**D**) Schematic of AT2 cells of transitional substates toward AT1 cells correlating with AP-1 activity (pink). Created with BioRender.com. (**E**) Heatmaps of top 25 T2 genes (Log_2_FC > 1, pval < 0.05; top) and their top associated peaks (Log_2_FC > 1, pval <0.05; bottom) that contain AP-1 motifs. (**F**) Top 30 gene ontology terms associated with T2 module genes. (**G**) Coverage plot of the top T2 module score gene *Cldn4* associated with significantly gained peaks that contain one or more AP-1 motifs (gray highlight). (Scale bar 50µm)

The emergence and shift of substates were asynchronous as they coexisted at 14 dpi. T2 formed small clusters separate from other AT2 cells and were limited to what we previously described as AT2-less regions and in-situ repair ^27,31^ (Fig. 3C, Fig. S2). Besides AT2-less regions, T1, T3, and T4 were also found in regions with normal-looking alveolar lumens and abundant lineage-traced, intermixed AT1 and AT2 cells, previously posited as the result of compensatory de novo growth ^27,32^ (Fig. S2). Congruently, KRT8^high^ activated AT2 cells in de novo regions lacked CLDN4 (Fig. S2A).

To delve into the T2 substate and its AP-1 connection, we linked single-cell multiome peaks to their nearest genes, defining T2-specific RNA and chromatin modules by selecting genes whose upregulation matched increased accessibility at AP-1–containing peaks (Fig. 3E). The resulting T2 RNA module included canonical transitional markers such as *Cldn4* and *Itga2*, and was enriched for gene ontology terms associated with cell adhesion, TGFβ signaling, wound healing, and inflammation—key features of transitional AT2 cells ^12,15,17^ (Fig. 3F). At the *Cldn4* locus, three AP-1–containing peaks showed maximal accessibility alongside the highest RNA expression in T2 (Fig. 3G). Although other injured AT2 subsets and their AT1 progeny showed slight increases in *Cldn4* RNA and chromatin accessibility, these lower-level signals were attributed to AP-1 activation spilling over to parallel processes, such as AT2 cell proliferation, and AT1 differentiation (Fig. 3G).

Collectively, our sc-multiome and immunostaining delineate diverse AT2 populations in Sendai virus–injured lungs, with AP-1 activation peaking in the CLDN4⁺ T2 substate.

### AP-1 members FOS, FOSB, and JUNB are required for CLDN4 expression and AT2 cell dispersion

Given AP-1’s diverse, context-dependent roles in lung tumorigenesis, fibroblast activation, and skin wound repair ^33–35^, we immunostained for all seven AP-1 family members (FOS, FOSB, FOSL1, FOSL2, JUN, JUNB, and JUND) in lungs harvested at 14 dpi with Sendai virus (Fig. 4A, 4E, and S3). While FOSL1, FOSL2, and JUND were absent, and JUN and the GFP-tagged *Fos* allele were constitutively present, only FOSB and JUNB specifically localized to *Sftpc^CreER^* lineage–traced AT2 cells after infection, but not in mock-treated controls. Across approximately 4,100 cells from 3 mice, 7-16% of AT2 cells upregulated FOSB and JUNB, and 30–40% of these cells lost LAMP3 expression, consistent with entry into the transitional state (Fig. 4A, 4B). Airway cells at baseline and airway-derived KRT5⁺ pods expressed FOSL2, consistent with its reported role in airway cells ^36^ (Fig. S3B).

**Fig. 4.**
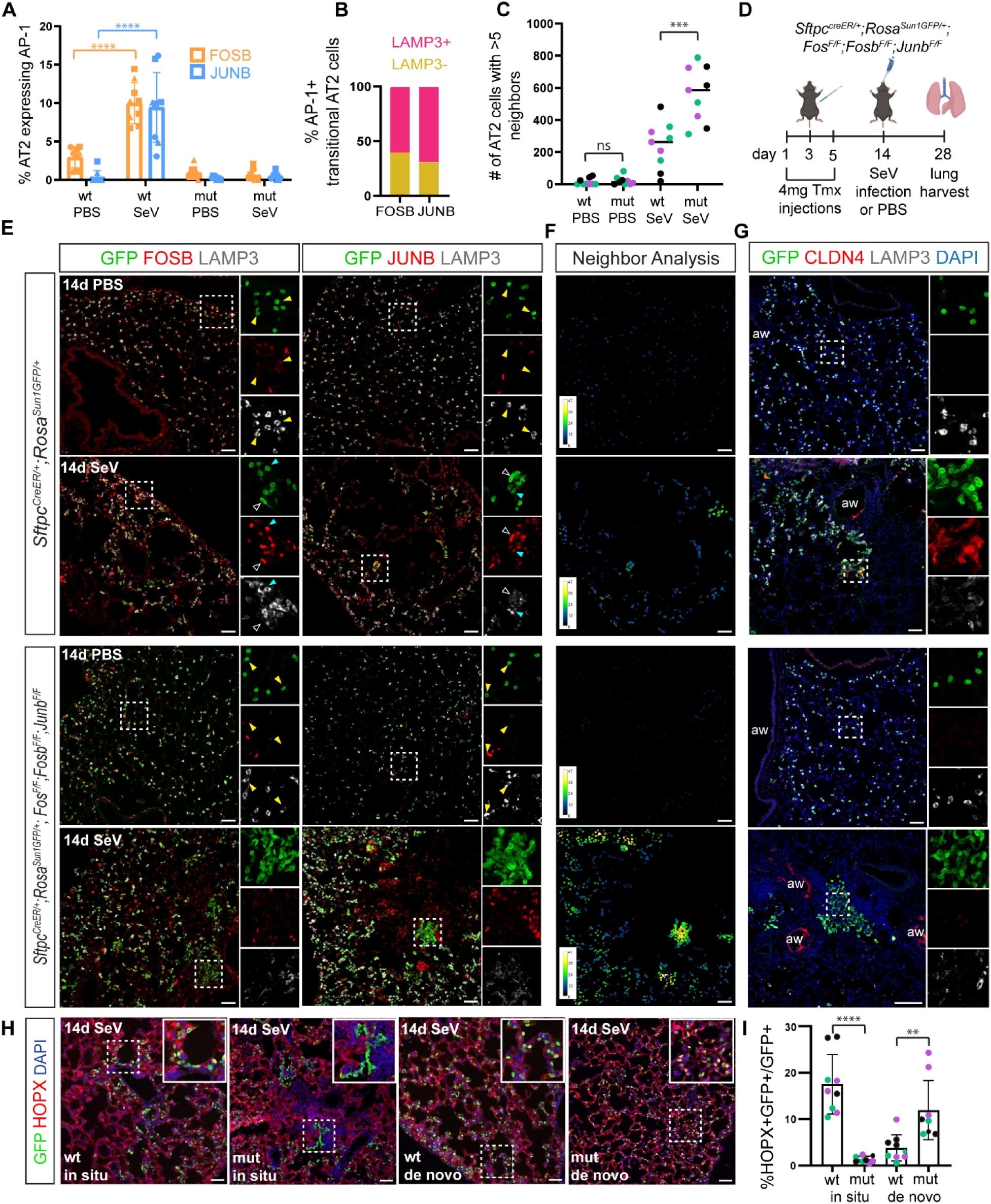
AP-1 members FOS, FOSB, and JUNB are required for CLDN4 expression, AT2 cell dispersion, and region-specific AT1 differentiation. (**A**) Quantification of AT2 cells upregulating FOSB and JUNB 14 days post SeV infection and their expected loss in the AP-1 mutant (mut). Squares, triangles, and circles represent 3 replicate mice (unpaired t-test, p-value < 0.0001). wt, wildtype. (**B**) Quantification of the percentage of FOSB or JUNB-expressing AT2 lineage-labeled cells that have lost LAMP3 expression, indicative of a transitional state. (**C**) ImageJ BioVoxxel neighbor analysis of AT2 cells with >5 AT2 neighbors showing increased clustering in the infected mutant. Colors representative of 3 replicate mice (unpaired t-test, p-value = 0.0006). (**D**) Experimental timeline of the AT2-specific AP-1 mutant deleting *Fos*, *Fosb*, and *Junb*. (**E**) Confocal images of 10µm sectioned lungs from PBS or SeV infected wildtype or AT2-specific AP1-mutant mice, showing upregulation of FOSB and JUNB (red) in AT2 cells (GFP) and partial loss of AT2 identity (LAMP3) upon infection in the wildtype (top panel). The AP-1 mutant lose FOSB and JUNB and have increased cell clustering with loss of AT2 identity (LAMP3) (bottom panel). Yellow arrowheads indicate AT2 cells that are LAMP3+ and do not express FOSB or JUNB, representative of a resting state. Cyan arrows indicate AT2 cells that are LAMP3+ and express FOSB or JUNB. Hollow arrows indicate AT2 cells that have lost LAMP3 expression while expressing FOSB or JUNB, representative of an AP-1+ transitional cell state. (**F**) ImageJ BioVoxxel neighbor analysis of the adjacent JUNB immunostaining panel, color-coded for AT2 neighbor counts ranging from 0 to 49. (**G**) Confocal images of 10µm sectioned lungs from AT2 lineage traced wildtype and AP-1 mutant mice showing a lack of infection-induced CLDN4 expression in mutant AT2 cells, which have lost LAMP3 expression and entered a transitional state. aw, airway. (**H**) Confocal images of 10µm sectioned lungs of SeV infected AT2 lineage traced wildtype or AP-1 mutant mice within AT2-less regions of damage (in situ) and intact, uninjured regions (de novo). (**I**) Quantification of HOPX+ AT2 cells within in situ (unpaired t-test, p-value < 0.0001) and de novo (unpaired t-test, p-value = 0.003) regions of wildtype and mutant mice. Colors represent 3 replicate mice. (Scale bar 50um).

Anticipating functional redundancy among AP-1 family members ^37,38^, we generated AT2-specific triple knockouts of *Fos*, *Fosb*, and *Junb* and induced recombination two weeks prior to Sendai virus infection (Fig. 4D). Deletion of induced FOSB and JUNB was confirmed by immunostaining (Fig. 4A, 4E). Recombination of the *Fos* allele was confirmed by widespread cytoplasmic GFP expressed upon recombination from the engineered *Fos* allele—distinct from the stronger, nuclear *Rosa^Sun1GFP^* reporter (Fig. S3C, 4E)—and by subsequent single-cell multiome reads mapping to its unique engineered 3′ UTR and verifying excision of the coding sequence.

The most striking phenotype in *Fos/Fosb/Junb* mutant lungs was the emergence of extensive AT2 cell clusters (Fig. 4C, 4E, 4F). In mock-infected lungs, AT2 cells in both wildtype and mutant lungs had 2-4 neighbors within a 50 µm radius; following infection, wildtype AT2 cells often grouped with >5 neighbors—consistent with proliferation—whereas mutant cells were more densely clustered with up to 50 neighbors, indicating impaired dispersion of daughter cells as proliferation remained unchanged (Fig. S4).

Intriguingly, these mutant clusters showed low LAMP3 but lacked CLDN4 expression, signifying entry into a transitional state without adopting the CLDN4⁺ T2 substate (Fig. 4G). Neither were they progressing toward T3, T4 or AT1 states as they were negative for HOPX (Fig. 4H). However, mutant cells in de novo regions did not over-cluster and instead were more frequently HOPX+ than their counterpart of the wildtype lung, perhaps compensating for the defective transition in the in-situ regions (Fig. 4H, 4I).

Thus, injury-induced FOSB and JUNB, together with constitutive FOS, are essential for promoting AT2 cell dispersion, the T2 marker CLDN4, and progression toward AT1 cells in in-situ regions.

### AT2-specific Fos/Fosb/Junb deletion blunts the CLDN4+ substate and attenuates fibroblast-activating senescence signals

To dissect genome-wide roles of FOS, FOSB, and JUNB, we performed single-cell multiome sequencing on sorted epithelial cells (CDH1+) from AT2-specific triple mutant lungs 14 days after mock or Sendai virus infection (Fig. 5A). Integration with the wildtype atlas (Fig. 2A) identified equivalent populations in the mutant, albeit with a downward shift of the T4 cluster in the UMAP (T4wt vs T4m; Fig. 5A). Transitional substates were also labelled in mock-infected wildtype and mutant lungs likely from data integration, but they lacked *Cldn4* expression and AP-1 motif accessibility and comprised 3% of AT2 cells, versus 12% in infected wildtypes (Fig. 5A, 5C, 5E). In infected mutants, the transitional fraction fell to 6%, with diminished T2 RNA module score including *Cldn4* expression as well as AP-1 motif (Fig. 5). The residual T2 population, not the broader AT2 cohort, retained significant *Fos* transcripts—suggesting that Cre-escaper cells preferentially adopted the T2 substate (Fig. 5B), though redundancy with and/or compensation by other AP-1 family members remained possible ^37,38^.

**Fig. 5.**
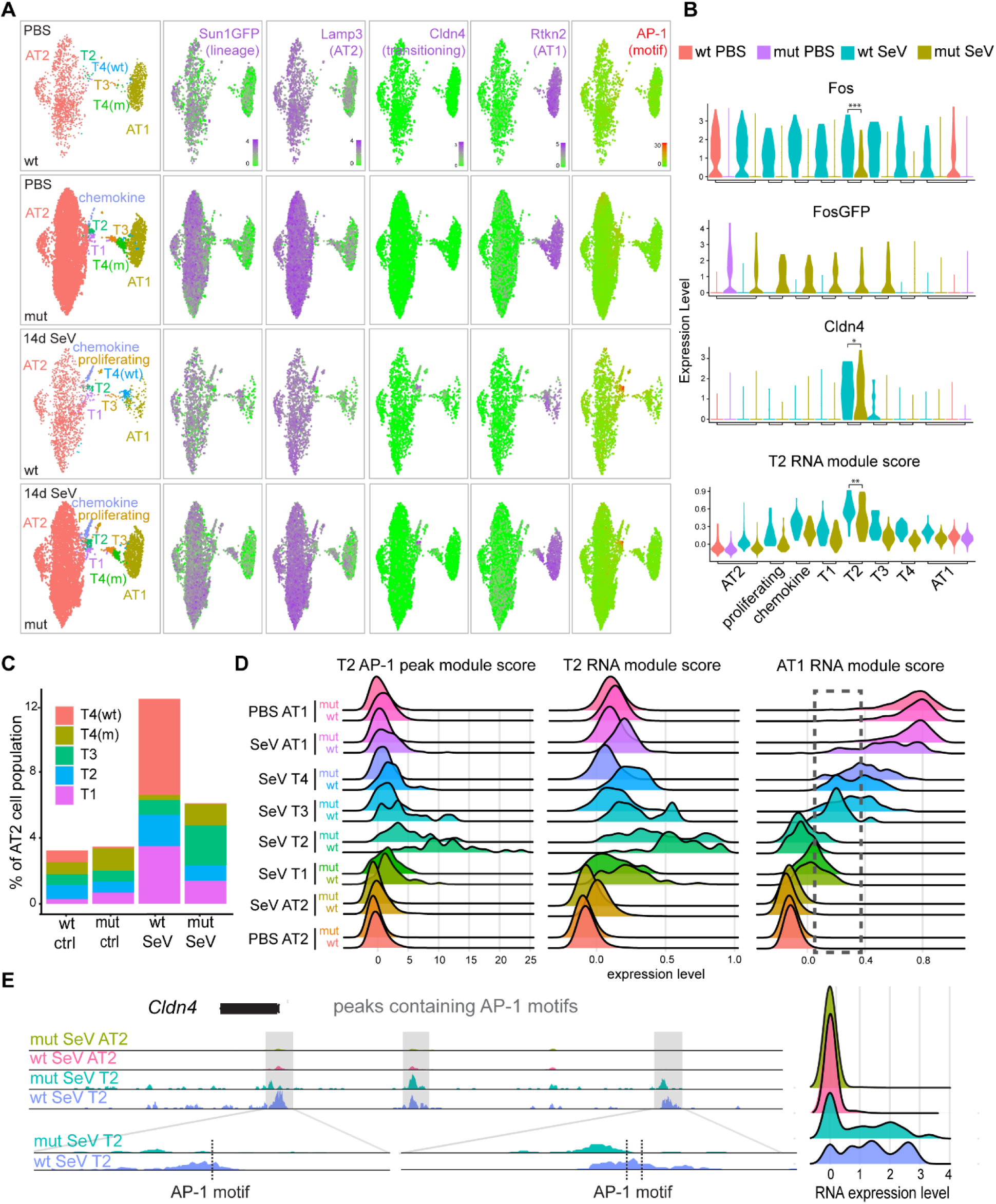
AT2-specific *Fos/Fosb/Junb* deletion blunts the CLDN4+ substate and enhances overall AT1 differentiation. (**A**) Single cell multiome UMAPs and feature plots of PBS and SeV infected *Sftpc^CreER/+^;Rosa^Sun1GFP/+^*;Fos*^F/F^*;Fosb*^F/F^*;Junb*^F/F^* mouse lungs integrated with the wildtype multiomic dataset (Fig. 2A). Feature plots show decreased AP-1 motif accessibility in the T2 population of the AP-1 mutant mice. (**B**) Single-cell multiome violin plots showing likely Cre-escapers in the mutant with residual *Fos* (unpaired Wilcoxon ranked sum test, p-value = 0.0001) expression preferentially in T2, and consequently a reduced but persistent level of *Cldn4* (unpaired Wilcoxon ranked sum test, p-value = 0.042) and T2 RNA module score (unpaired Wilcoxon ranked sum test, p-value = 0.001). The post-recombination *FosGFP* transcript is limited to AT2 cells of the AP-1 mutant. (**C**) Single-cell multiome stacked barplot of the percentages of transitional cell states within all AT2 cells for each condition. (**D**) Single-cell multiome ridgeplots of T2 RNA and ATAC module scores showing leftward (lower) distribution shifts across mutant populations, as well as ridgeplot of AT1 RNA score. Mutant T3, T4, and AT1 populations have an increased AT1 module expression over their wildtype counterparts. (**E**) Single-cell multiome coverage plots (left) of 3 *Cldn4* associated peaks containing AP-1 motifs (gray highlight and dotted lines). Two out of 3 peaks are reduced or shifted in the mutant associated with reduced *Cldn4* expression shown in the color-coded RNA ridgeplot (right).

To capture both genetic and phenotypic heterogeneity across Seurat clusters, we generated ridge plots comparing T2-specific chromatin and RNA module scores in wildtype versus AP-1 triple-mutant cells (Fig. 5D). As anticipated, the T2 chromatin module was restricted to the T2 population in wildtype but became negligible in mutants. For instance, two of the three AP-1–motif–containing peaks at the *Cldn4* locus were lost or shifted upon *Fos/Fosb/Junb* deletion (Fig. 5E). By comparison, the T2 RNA module—which included *Cldn4*—was also detectable at lower levels in T1, T3, T4, and even AT1 cells in wildtype lungs, likely reflecting RNA perdurance during substate transitions or parallel injury programs sharing genes. In the mutant, this RNA signature was reduced yet remained detectable in T1 and T2 populations, but negligible in T3 and T4 populations (Fig. 5D). AT1 genes were upregulated in mutant T3/T4 and AT1 clusters that encompassed those derived from AT2 cells (Fig. 5D). This rightward shift was still observed within *Rosa^Sun1GFP^* lineage-traced cells (Fig. S5), presumably dominated by the higher abundance of HOPX⁺ lineage-traced cells in de novo regions (Fig. 4H, 4I).

To assess the non–cell-autonomous consequences of AP-1 loss in AT2 cells, we analyzed the senescence-associated secretory phenotype (SASP) in transitional AT2 cells known to activate fibroblasts ^18,39^. Using a KEGG-derived SASP RNA module score, we observed that Sendai virus injury increased SASP expression in all substates—peaking in T2—but this induction was attenuated in AP-1 mutants, particularly in T4 where SASP normally subsided, although chromatin peaks linked to SASP genes showed no accessibility changes upon *Fos/Fosb/Junb* deletion (Fig. 6A). Scatterplots comparing injury-induced and AP-1–dependent SASP genes highlighted the Cdk, Mapk, and Tgfb families (Fig. 6B). Congruent with these data, the fibrosis marker SFRP1 was induced in fibroblasts adjacent to transitional AT2 cells with diminished LAMP3 after infection; in AP-1 mutants, SFRP1+ cells were rare and possibly limited to escaper AT2 cells (Fig. 6C). By comparison, both wildtype and AP-1 mutant lungs had minimal SFRP1 expression in de novo regions with similarly diminished LAMP3 albeit lack of CLDN4 expression (Fig. S2), but high SFRP1 expression near airways from which KRT5+ pods were known to spread into the alveolar region, possibly induced by an airway SASP program spared by *Sftpc^CreER^* (Fig. 6C).

**Fig. 6.**
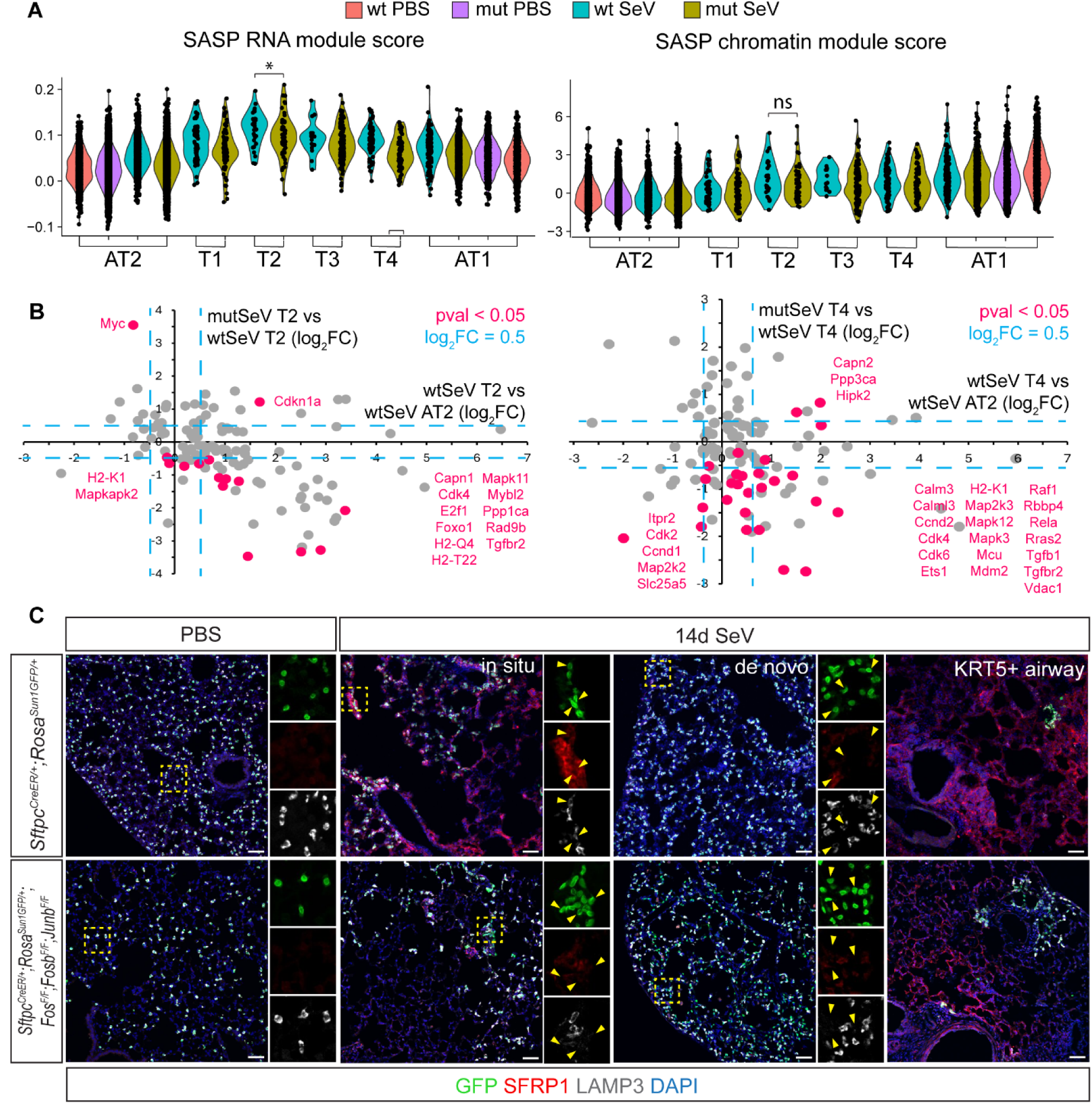
AT2-specific *Fos/Fosb/Junb* deletion attenuates fibroblast-activating senescence signals. (**A**) Single-cell multiome violin plots of KEGG-generated SASP RNA and chromatin module scores showing lowered RNA module scores in the AP-1 mutant populations but comparable chromatin accessibility between wildtype and mutant (unpaired Wilcoxon ranked sum test, p-value = 0.044). (**B**) Scatterplots of log_2_FC SASP gene expression. X axes show infection-induced upregulation of highlighted significant genes in the T2 (left) or T4 (right) substate compared to AT2 cells in the wildtype; Y axes show their downregulation due to AP-1 mutation. (**C**) Confocal images of 10µm sectioned lungs showing induction of a fibrotic mesenchymal marker SFRP1 selectively in AT2-less regions (in situ) adjacent to transitioning AT2 cells (GFP+LAMP3-) in infected wildtype lungs but not AP-1 mutant lungs. Both wildtype and AP-1 mutant lungs have weak SFRP1 in de novo regions and strong SFRP1 near KRT5+ airway pods. (Scale bar 50um).

Collectively, our findings demonstrate that during lung injury-repair, AT2 cell–expressed FOS, FOSB, and JUNB drive the CLDN4⁺ T2 substate—enhancing SASP-mediated fibroblast activation.

### AP-1 activation marks oncogenic transition in murine models and human lung adenocarcinoma

We reasoned that AT2 cells might also undergo activation, potentially permanent, during tumorigenesis, particularly given our prior report of KRT8+ alveolar cells (KACs) in human and mouse lung adenocarcinoma ^19^. To test this possibility, we used an AT2-specific oncogenic Kras expression mouse model. Fourteen days after Cre activation, *Rosa^tdT^* lineage-traced AT2 cells were purified from wildtype or *Kras^LSLG12D^* lungs and analyzed by bulk ATAC-seq (Fig. 7A). In *Kras^LSLG12D^* lungs, a set of 17,621 peaks gained accessibility and were enriched for AP-1 motifs along with GATA and TEAD motifs, whereas another set of 8,509 peaks lost accessibility and were enriched for NKX and FOXA motifs (Fig. 7B). Notably, the same three AP-1 motif–containing peaks at the *Cldn4* locus that were reversibly opened during Sendai infection became accessible in *Kras* mutant AT2 cells (Fig. 7C).

**Fig. 7.**
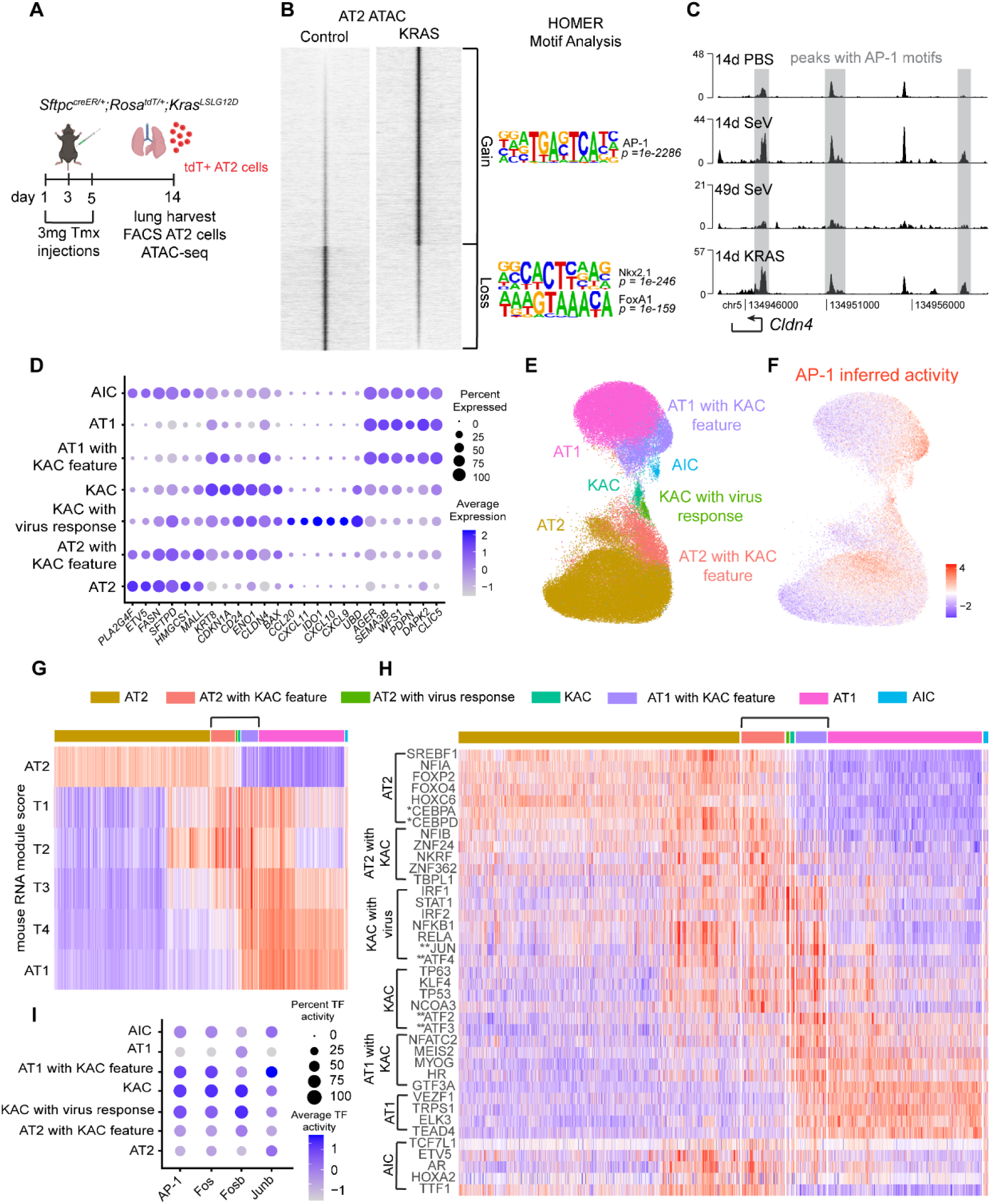
AP-1 activation marks oncogenic transition in murine models and human lung adenocarcinoma. (**A**) Schematic of AT2 cell lineage labeling, induction of oncogenic *Kras*, and lung harvest for bulk ATAC-seq. (**B**) Heatmap of bulk ATAC-seq gained peaks (Log_2_FC > 1, pval < 0.05) and lost peaks (log_2_FC < −1, pval < 0.05) from KRAS vs. control with top HOMER motifs. (**C**) Coverage plot of *Cldn4* in AT2 cells from PBS treated, 14 and 49 days post Sendai infection, and 14 days post oncogenic *Kras* induction lungs. Gray bars highlight peaks that contain one or more AP-1 motifs. Plot axes were normalized based on fraction of reads in each peak (FriP). (**D**) Single-cell RNA-seq dotplot of top 6 AT2, KAC (*KRT8+* alveolar intermediate cells), virus response, and AT1 genes identifying each cluster in human lung adenocarcinoma (LUAD). AIC: alveolar intermediate cell with both AT1 and AT2 markers. (**E**) UMAP of alveolar cells from human LUAD. (**F**) UMAP of inferred AP-1 transcription factor activity amongst alveolar human LUAD populations. (**G**) Heatmap of RNA module scores from the T1-T4 populations of Sendai infected mice in the human LUAD populations. The T2 score is the highest in KAC-related populations. Bracket indicates populations with a KAC response. (**H**) Heatmap of top 5 inferred transcription factor activity for each human LUAD population. *, transcription factors known to be active within AT2 cells and found within the top 10 hits for inferred transcription factor activity within the AT2 cell population. **, AP-1 members found within the top 50 hits of each cluster. Bracket indicates populations with a KAC response. (**I**) Dotplot of inferred transcription factor activity of AP-1 members Fos, Fosb, and Junb showing highest activation within KACs.

To assess the relevance to human lung adenocarcinoma, we analyzed our snRNA-seq dataset of normal lung tissues, precursor lesions and invasive adenocarcinoma, comprising 83,790 alveolar epithelial cells from 23 patients (in press). In addition to KACs (*KRT8*+ alveolar intermediate cells) defined by *KRT8*, we identified AT1 and AT2 subsets carrying the KAC gene signature that we previously derived, a KAC subset enriched for virus response genes such as *CXCL9* and *CXCL10*, and an alveolar intermediate cell (AIC) population characterized by *AGER* and *SFTPD* (Fig. 7D, 7E). To compare this cancer-associated heterogeneity with that observed during mouse Sendai virus infection, we scored human cell populations using orthologous gene modules derived from mouse populations (Fig. 7G). KACs showed the strongest alignment with T1 and T2 substates, whereas T3 and T4 substates increasingly aligned with human AT1 cells, as they did with mouse AT1 cells (Fig. 7G). In the absence of ATAC-seq data from human specimens, we inferred transcription factor (TF) activities by using decoupleR, which computes TF activity scores based on a network of TFs, their targets, and weights of the TF-target interaction (Fig. 7F, 7H, 7I)^40^. AP-1 activity was highest in KACs, including those with virus responses, as well as in their variants among AT1 and AT2 cells. Identification of top transcription factor activities per cluster confirmed known transcription factors for AT2 cells (CEBP and FOX) and revealed AP-1 transcription factors within the top 50 hits of the KAC and KAC with virus response populations (Fig. 7H). Of the three AP-1 members deleted within the Sendai infected mice, Fos and Fosb showed highest transcription factor activity in the KAC and KAC with virus response clusters, supporting a conserved process of AP-1–mediated AT2 cell activation (Fig. 7I).

## DISCUSSION

Our integrated analysis of RNA expression and chromatin accessibility refines transitional AT2 cells into four substates, corroborated by immunostaining. The T2 substate features CLDN4 and AP-1 motifs and requires constitutive FOS and induced FOSB/JUNB for AT2 cell dispersion, fibroblast signaling, and AT1 differentiation. AP-1 activation is conserved in mouse AT2 cells carrying oncogenic Kras and in transitional AT2 cells from tissues representing the pathologic ontology of early-stage human lung adenocarcinoma. Our enhanced understanding of transitional AT2 cell heterogeneity and key transcriptional regulators provides a foundation for dissecting the complex gene regulatory networks involved and highlights potential roles of AP-1 in AT2 cells in lung fibrosis and cancer.

Transitional AT2 cells have been documented in numerous injury models and human diseases and have been proposed as either a cause or a consequence of tissue restructuring, with both detrimental and beneficial effects ^41,42^. For example, *Ctgf*+ versus *Lgals3*+ and primed versus damage-associated AT2 cells have been described in the mouse lung, while various aberrant basaloid intermediates and KRT8+ alveolar cells have been detected in the human lung ^15,17,19^. Such phenotypic heterogeneity may stem from the functional heterogeneity of AT2 cells—as specialized surfactant producers, barrier formers, and plastic progenitors capable of differentiating into AT1 cells, as elaborated below.

First, the CLDN4+ T2 substate defined in our study shows reduced AT2 and AT1 gene signatures (Fig. 2B), diverting cellular resources away from gas exchange physiology. Second, T2’s defining feature of AP-1 activation—typically associated with immediate early responses ^43^—along with its signature gene ontology terms for cell adhesion and migration (Fig. 3F) and AP-1–dependent fibroblast-activating signals (Fig. 6A, 6B), supports an emergent role in barrier restoration. Such barrier restoration is likely an ancient program across external and mucosal surfaces ^44,45^ but, in the lung alveoli, is supplemented by newly differentiated AT1 cells and even invading airway p63+ cells. Intriguingly, AT2-less regions following Sendai infection, which presumably have breached barriers, require in situ repair that triggers CLDN4+ AT2 cells and SFRP1+ fibroblasts (Fig. 6C). By contrast, otherwise normal regions with reduced LAMP3 but minimal CLDN4 and SFRP1 expression likely represent de novo growth to compensate for virus-damaged gas exchange surface area. Consistently, compensatory growth after pneumonectomy also barely induces SFN+ transitional cells and SFRP1+ fibroblasts ^39^. Third, AT2 differentiation into AT1 cells also exhibits regional differences. In in situ regions, AP-1 mutant AT2 cells downregulate LAMP3 but fail to efficiently activate CLDN4 or differentiate into AT1 cells, implicating the T2 substate as an intermediate toward AT1 differentiation. In contrast, AP-1 mutant AT2 cells in de novo regions differentiate into AT1 cells more robustly, suggesting a T2-independent differentiation pathway. The intertwining of region-specific barrier restoration and compensatory growth is also likely relevant in lung fibrosis and cancer, highlighting the need for precision analyses to isolate specific components and regulators.

Parallel programs in AT2 cells also unfold over time. Most notably, an interferon response to contain the Sendai virus peaks at 7 days post infection across epithelial, endothelial, and likely other lineages, and persists beyond day 14 when tissue repair is already under way ^27,46^. Residual inflammation in AT2 cells may be orthogonal to their transitional states, influencing them only indirectly by diverting cellular machinery such as protein synthesis and metabolism at the individual cell level, or by shaping the extent of viral injury at the tissue level. Nevertheless, memory of prior events recorded in the epigenome, cellular structure, and composition has been described in the skin, coincidentally also mediated by AP-1 ^47^. Future studies are needed to identify the specific functional AP-1 members in AT2 cells, assess their long-term effects on subsequent injuries or tumorigenesis, and define their intersection with other known transcriptional regulators such as p53 and ATF stress factors ^48–50^.

## METHODS

### Mice

The following mouse strains were used: *Sftpc^CreER 51^*, *Rosa^tdT^* ^52^, *Rosa^Sun1GFP^* ^53^, and *Fos^F/F^;Fosb^F/F^;Junb^F/F^* ^54^, *Kras^G12D^* ^55^. Tamoxifen (T5649, Sigma) was dissolved in corn oil (C8267, Sigma) and administered through intraperitoneal injections at doses, timepoints, and sample size explained in figure legends. Power analysis was not used to calculate sample sizes. Experimental mice were of mixed genders and mixed backgrounds of C57BL/6 and 129. Mice were housed at 22 °C 45% humidity, and 12-12-h light-dark cycle conditions within the MD Anderson facility. All animal experiments were approved by the Institutional Animal Care and Use Committee at the University of Texas MD Anderson Cancer Center.

### SeV Infection

Mice were infected following our previously published protocol ^27^. Mice 8 weeks of age were briefly anesthetized with isoflurane and suspended by their upper incisors with a 60° incline. A non-lethal dose of approximately 2.1 × 10^7^ plaque-forming units of Sendai Virus (ATCC #VR-105, RRID:SCR_001672CSCSSCS) diluted in 40 µL of PBS was administered through oropharyngeal installation. Control groups received 40 µL of PBS.

### Section Immunostaining

Mice were first given an intraperitoneal injection of Avertin (Sigma, T48402). To flush blood out of the lungs, the heart was perfused through the right ventricle with PBS. Lungs were inflated through a cannulated trachea with 0.5% paraformaldehyde (Sigma, P6148) in PBS at a pressure of 25 cm H_2_O. Lungs were removed and fixed in 0.5% PFA for 3-6 hours at room temperature. Lungs were washed with PBS to remove fixative and left overnight at 4°C in PBS. For cryopreservation, lung lobes were carefully separated using forceps and incubated overnight in 20% sucrose with 10% optimal cutting temperature compound (OCT; 4583, Tissue-Tek, Tokyo, Japan) in PBS for cryoprotection. Lobes were embedded into blocks with OCT and frozen. Tissue blocks were sectioned at a thickness of 10 µm and tissue was blocked for 1 hour in PBST (0.3% Triton X-100 in PBS) with 5% normal donkey serum (017-000-121, Jackson ImmunoResearch). Primary antibodies were diluted in PBST and added to the tissue for overnight incubation in a humidified chamber at 4°C. Slides were then washed in PBS for 1 hour at room temperature. Secondary antibodies (Jackson ImmunoResearch) and 4′,6-diamidino-2-phenylindole (DAPI) were diluted in PBST and incubated with the sections for 1 hour at room temperature. Tissue sections were washed in PBS as done previously, and mounted using Aquapolymount (18606, Polysciences). Stained tissue sections were imaged using a confocal microscope (Olympus FV1000) and analyzed with FIJI/ImageJ (version 2.16.0).

### Antibodies

The following antibodies were used for section immunostaining: rat TROMA-I anti-cytokeratin 8 (KRT8, 1:5000, University of Iowa Hybridoma Bank), guinea pig LAMP3 antibody (LAMP3, 1:500, 391005, Synaptic Systems), mouse Hop Antibody (E-1) AF647 (HOPX, 1:500, sc-398703, Santa Cruz Biotechnology), rabbit Claudin 4 polyclonal antibody (CLDN4, 1:250, 36-4800, Thermo Fischer Scientific), purified anti-mouse I-A/I-E Antibody (MCHII, 1:500, 107601, Biolegend), chicken anti-GFP antibody (GFP, 1:250, AB13970, Abcam), FosB (5G4) rabbit mAb (FOSB, 1:250, 2251, Cell Signaling), JunB (C37F9) rabbit mAb (JUNB, 1:250, 3753, Cell Signaling), goat FRA1 antibody (FOSL1, 1:250, AF4935 R&D Systems, rat anti-FRA-2 antibody clone REY146C (FOSL2, 1:250, MABS1261, Millipore Sigma), rabbit anti-JUND antibody (JUND, 1:250, SAB4501615, Millipore Sigma), rabbit Cytokeratin 5 (EP1601Y) monoclonal primary antibody (KRT5, 1:500, 305R-15, Sigma), rabbit Ki-67 recombinant monoclonal antibody (SP6) (MKI67, 1:1000, MA5-14520, ThermoFischer Scientific), rabbit anti-SFRP1 antibody (SFRP1, 1:250, ab126613, Abcam).

### Cell dissociation and FACS

Collection and dissociation of mouse lungs for FACS and single cell multiome sequencing preparation have been previously defined in our published procedure ^32^ with minor modifications. Lung lobes were separated and finely minced with forceps. Accessory lobes were saved for cryopreservation and section immunostaining. Lungs were digested for 45 minutes at 37°C in the following solution: 1.35 mL Leibovitz media (Gibco, 21083-027), 2 mg/mL collagenase type I (Worthington, CLS-1, LS004197), 0.5 mg/ml DNase I (Worthington, D, LS002007), and 2 mg/ml elastase (Worthingon, ESL, LS002294). Digest was carefully mixed halfway through digestion to aid with tissue dissociation. Enzymatic activity was stopped with the addition of 300 µL of fetal bovine serum (FBS, Invitrogen, 10082-139). Tissue was carefully mixed with a pipette to dissociate into single cells and filtered through a 70 µm strainer (Falcon, 352350). For single cell multiome experiments, each condition had n=2 mice (1 male and 1 female) that were digested separately for ease of dissociation and mixed together at this step. Contents were then transferred to a 2 mL tube on ice and the remaining steps were conducted in a 4°C cold room. The cells were centrifuged at 1537 rcf for 1 minute and the supernatant was removed. The cells were then resuspended in 1mL of red blood cell lysis buffer (15 mM NH_4_Cl, 12 mM NaHCO3, 0.1 mM EDTA, pH 8.0) and incubated on ice for 3 minutes. This step was repeated to make sure all red blood cells were depleted and a white pellet was observed. Samples were washed with 1mL of Leibovitz with 10% FBS and resuspended again in 1mL of Leibovitz with 10% FBS. For single cell multiome sequencing, cells were stained with the following antibodies at 1:250 concentration for 30 minutes on ice: CD45-PE/Cy7 (BioLegend, 103114), ECAD-PE (BioLegend, 147304), and ICAM2-A647 (Invitrogen, A15452). SYTOX blue (Invitrogen, S34857) was added at a concentration of 1:1000 and the samples were run through a 35 µm filter into a 5 mL FACS tube just before sorting using an AriaII cell sorter with a 70 µm nozzle at 4°C. ECAD positive epithelial cells were collected from the subsequent negative gating of the CD45 and ICAM2 populations. Single cell libraries were generated using the Single Cell Multiome ATAC + Gene Expression kit (10x Genomics) with10,000 nuclei were loaded per lane.

### Bulk ATAC seq and analysis

The sample preparation for bulk ATAC-sequencing was modified from the OMNI-ATAC sequencing protocol ^56^ and previously described ^57^. Sendai infected and control *Sftpc^CreER/^*+;Rosa*^tdT/+^* mouse lungs were harvested, dissociated and sorted for 100,000 tdT+ AT2 cells. Cells were centrifuged in 4°C at 1098 rcf for 5 minutes in a fixed angle centrifuge. The supernatant was discarded, and the pellets were resuspended in 50 µL of cold ATAC-RSB lysis buffer (0.1% NP-40, 0.1% Tween-20, 0.01% Digitonin, 10 mM Tris-HCl pH 8.1, 10 mM NaCl, 3 mM MgCl_2_). The resuspension was incubated on ice for 3 minutes followed by the addition of 1mL of cold ATAC-RSB+Tween (10 mM Tris-HCl pH 8.1, 10 mM NaCl, 3 mM MgCl_2_, 0.1% Tween-20) and inverted 3 times to mix. Nuclei were pelleted through centrifugation in 4°C at 1098 rcffor 10 minutes in a fixed angle centrifuge. The supernatant was discarded, and the pellet was resuspended in 50 µL of transposition buffer that included the following: 22.5 µL of Tn5 Reaction Mix (PBS, 2.2% Digitonin, 2.2% Tween-20), 25 µL of Tn5 Buffer (10 mM MgCl_2_, 20 mM Tris-HCl, 20% Dimethyl Formamide), and 2.5 µL of Tn5 Enzyme (NX#-TDE1, Tagment DNA Enzyme, 15027865, Illumina). The reaction was incubated in a thermocycler at 37°C for 30 minutes. DNA was purified using the Qiagen MinElute PCR Purification Kit (29004) following the manufacturer instructions with one modification of eluting with 10 µL of distilled water. Sample barcoding and PCR amplification was performed using the Greenleaf ATAC primers (^58^ following the OMNI-ATAC protocol PCR parameters. To assess amplification efficiency, 5 µL of each sample was saved and set aside while the remainder of each sample continued to size selection purification using SPRIselect beads (Beckman Coulter, B23318). Following the manufacturer instructions for double size selection, the sample was incubated for upper size selection with 0.5X the volume and for lower size selection at 1.8X the volume. Size selection was verified using agarose gel electrophoresis with 1 µL of final sample as well as the previously saved pre size selection sample. Final DNA quantification was measured using the Qubit HS dsDNA assay (Thermo Fisher Scientific, Q32851).

Samples were sequenced and processed as previously described (Little et al., 2021). To summarize, samples were sequenced on the Illumina Novaseq6000. Fastq files were generated through demultiplexing using BCL2Fastq and quality checked using FastQC (http://www.bioinformatics.babraham.ac.uk/projects/fastqc/). Poor quality bases were removed using Trimmomatic and filtered reads were aligned to the UCSC mm10 reference genome with Bowtie2. Aligned reads were converted from sam files to bam files using Samtools and filtered for poor alignment and unmapped reads using Picard, MarkDuplicates, and Samtools with the following parameters: –f 3 –F 4 –F 256 –F 1024 –F 2048 –q 30. Peak calling was conducted using MACS2 with the following parameters: -q 0.05 –nomodel –shift −100 –extsize 200 –broad and filtered for a−log10 q value greater than 5 and removal of mm10 blacklist sites. Differentially accessible regions were detected using Diffbind with a fixed peak width of 500bp between control and infected samples for normalization. Heat maps were generated using Easeq (^59^. Accessible motifs were identified using HOMER motif analysis with the following parameters: findMotifs.pl -mm10-size 200. HOMER AnnotatePeaks.pl was used to find genes associated with peaks containing AP-1 motifs. AP-1 motif files to run AnnotatePeaks.pl were downloaded from HOMER motif analysis outputs. Peaks with their associated genes from 14 day and 49 day post Sendai infection time points were matched and plotted against each other to generate a scatter plot using the MATCH function in Microsoft Excel. Peaks containing AP-1 motifs were matched and highlighted within the scatter plot.

Access to all bulk ATAC-seq data can be found on Gene Expression Omnibus repository under accession number GSE309751.

### Single-cell multiome seq and analysis

Cellranger-arc (version 2.0.0) was used to run “cellranger-arc count” on all samples to align fastq files to a custom mm10 reference genome containing the *Sun1GFP* and *Fos-GFP* transcripts. Wildtype control, wildtype Sendai infected, mutant control, and mutant Sendai infected samples were then aggregated together using “cellranger-arc aggr”. Outputs from Cellranger-arc were analyzed in R (version 4.3.2) using the Seurat (version 5.2.0) and Signac (version 1.14.0) packages for RNA and ATAC. To eliminate batch effect amongst the wildtype and mutant samples generated on different days, Harmony package ^60^ was used to integrate all datasets. Harmony was run on RNA and ATAC as separate reductions first and then combined as a reduction list in a weighted nearest neighbor analysis to generate umaps containing both modalities. AT2 and AT1 RNA module scores were generated from AT2 and AT1 gene lists previously published ^22,61^ using the AddModuleScore function in Seurat. Chromatin regions of AT1 and AT2 genes were generated using the LinkPeaks and GetLinkedPeaks functions in Signac to find peaks associated with the genes used in the RNA module scores. Once peaks were identified, the AddChromatinModule function was used to generate chromatin module scores for AT2 and AT1 cells. The same procedure was followed to produce AP-1 RNA and chromatin module scores using a list of all known AP-1 family genes ^62^. The Krt8 ADI ^12^ and PATS ^15^ gene lists were collected from published data.

### Image analysis

Image analysis was conducted using FIJI/ImageJ (version 2.16.0). AT2 cells expressing AP-1 were quantified using the Cell Counter plugin. Visualization and quantification of cell clustering was conducted using the Adjustable Watershed (https://imagej.net/plugins/adjustable-watershed/adjustable-watershed) and Biovoxxel 3D Box Neighbor Analysis 2D/3D (https://imagej.net/plugins/bv3dbox) plugins. First, the GFP+ channel was separated as its own image, switched to an 8-bit image, and thresholded to ensure all cells were visible without background. To help segregate clustered cells and identify them as an individual object, the Adjustable Watershed plugin was used at tolerance of 0.5. Cells were spot checked for accurate segregation by merging the watershed adjusted image back with the DAPI and GFP channels of the original image and made transparent to visualize if clustered cells were now truly separated. A line was hand drawn in between cells that were not successfully separated after using adjustable watershed to ensure that all cells represent individual objects. Biovoxxel Neighbor Analysis was then performed to generate intensity images of neighbor counts and quantifications of neighbors per object within 50 µm. Neighbor count intensity image scales were normalized to the image with highest intensity.

### Human single-nucleus RNA seq analysis

Single-nucleus RNA sequencing was performed using FFPE scrolls from 75 matched normal, precursor, and invasive lung adenocarcinoma samples from 23 patients including 24 normal, 24 lung adenocarcinoma (LUAD), 9 atypical adenomatous hyperplasia (AAH), 14 adenocarcinoma *in situ* (AIS), and 4 minimally invasive adenocarcinoma (MIA) tissue samples. Sequencing data was analyzed in R (version 4.4.1) using Seurat (version 5.3.0). Gene signatures obtained from AT1, AT2, and T1-T4 states from the mouse single-cell multiome sequencing data were converted to human orthologs using Gprofiler2^63^. Module scores for each population were generated in the Human dataset using the AddModuleScore function in Seruat. DecoupleR was used to infer transcription factor activity^40^. Differentially active TFs were identified using Wilcoxon’s rank-sum test by running the FindAllMarkers function on the activity scores obtained from DecoupleR.

Access to the human single-nucleus RNA seq data can be found on Gene Expression Omnibus repository under accession number GSE308103.

## Supporting information

mouse single-cell multiome seq R script

human single-cell RNA seq R script

## ACKNOWLEDGEMENTS

We thank Drs. Michael Greenberg at Harvard University for providing the *Fos/Fosb/Junb* mice, Harold Chapman at UCSF for providing the *Sftpc^CreER^* mice, and Antony Rodriguez at Baylor College of Medicine for providing *Sftpc^CreER/+^;Rosa^tdT/+^* mice. ChatGPT was used to edit the manuscript. This work was supported by the University of Texas MD Anderson Cancer Center Retention Fund (JC), the University of Texas MD Anderson Cancer Center Office of Strategic Research Programs (HK), National Institutes of Health grants R01HL130129, R01HL153511, R35HL171346 (JC) and F31HL165914 (AML), and the National Cancer Institute (NCI) grants U01CA264583 (HK) and R01CA272863 (HK).

## AUTHOR CONTRIBUTIONS

AML, HK, and JC designed research; AML, TN, SY and TZ performed research and analyzed data; AML, TN, SEE, HK, and JC wrote and edited the paper; all authors read and approved the paper.

---

**Table S1.** HOMER motif analysis results in Fig. 1

**Table S2.** Bulk ATAC-seq peaks in Fig. 1

**Table S3.** Bulk ATAC-seq scatter plot in Fig. 1

**Table S4.** Bulk ATAC-seq vs. pseudobulk single cell RNA-seq scatter plot in Fig. 1

**Table S5.** Single-cell multiome top 50 genes per cluster in Fig. 2

**Table S6.** Module score genes and peaks for AT2 cells, AT1, cells, PATS, K8 ADI, AP-1, and T1-T2 clusters (T2 genes with AP-1 peaks only and T2 all genes) in Fig. 2

**Table S7**. Single-cell multiome chromvar motif analysis per cluster in Fig. 2

**Table S8.** T2 RNA module score GO analysis in Fig. 3

**Table S9.** ImageJ quantification of AP-1 expression in AT2 cells bar and stacked bar graph in Fig. 4

**Table S10.** ImageJ AT2 clustering neighbor analysis raw data in Fig. 4

**Table S11.** ImageJ quantification of HOPX+ AT2 cells within in situ and de novo regions in Fig. 4

**Fig. S1.**
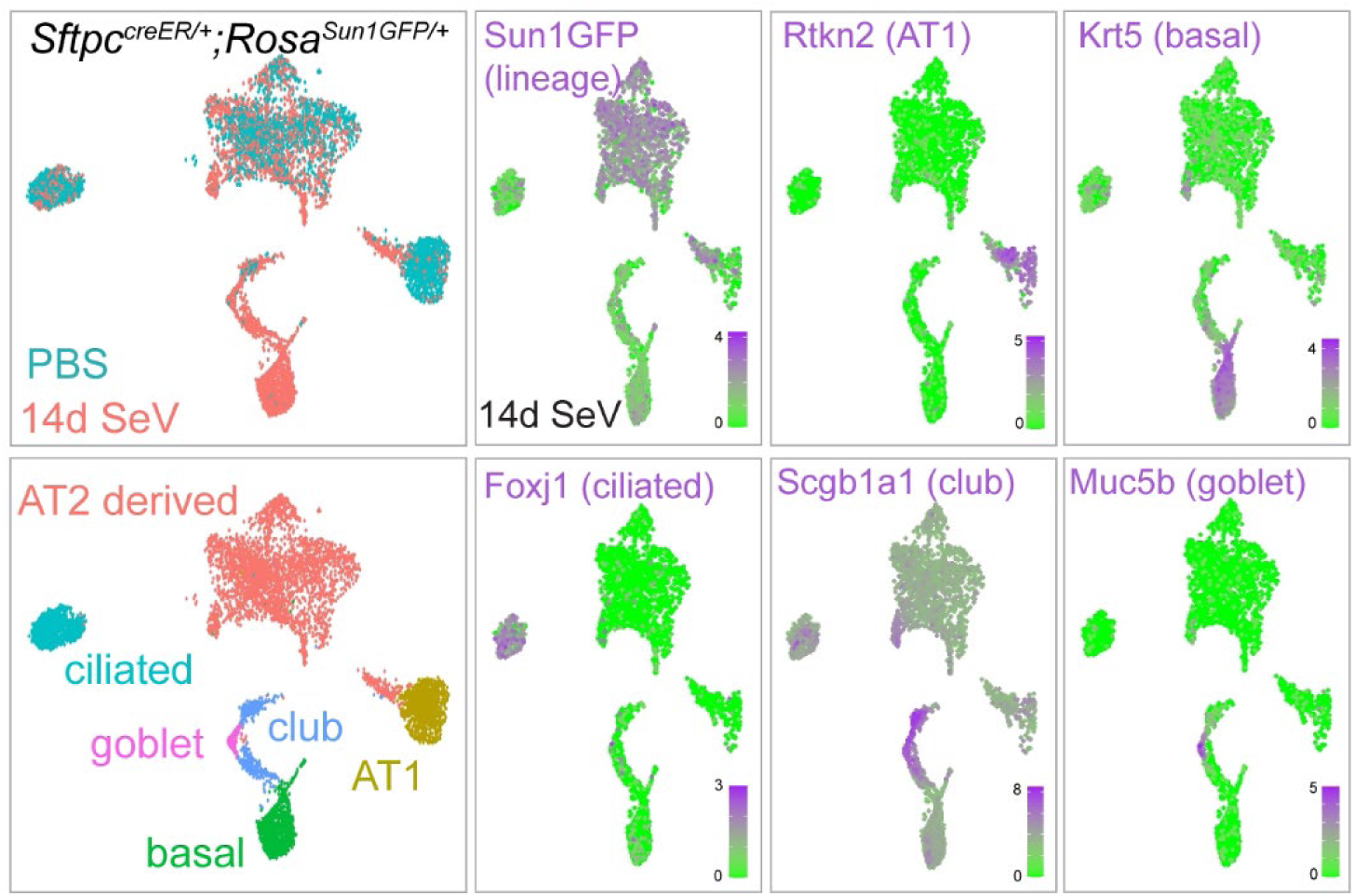
Single-cell multiome (ATAC+RNA) weighted nearest neighbor UMAPs of all epithelial cells. Dimplots color-coded for PBS control and SeV infected cells (top) and AT2, AT1 and 4 airway cell types (bottom). Feature plots of SeV infected UMAPs with representative gene expression from each cluster (purple). Expanded basal cells are infection-induced KRT5+ pods.

**Fig. S2.**
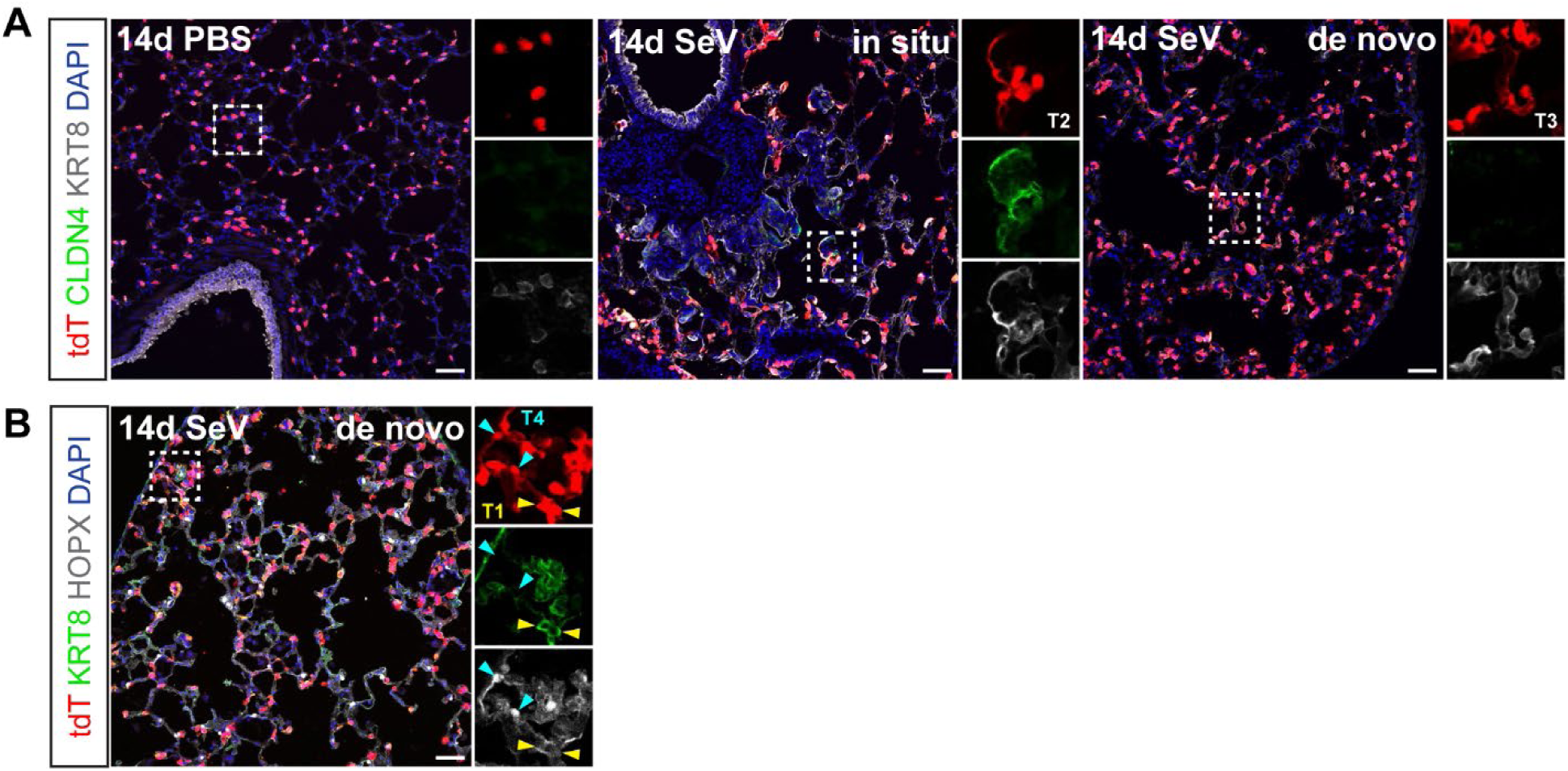
CLDN4+ T2 transitional cells are limited to in situ regions; T1/3/4 substates are also in de novo regions. Confocal images of 10µm sectioned lungs from PBS or SeV infected *Sftpc^CreER/+^;Rosa^tdT/+^* mice. (**A**) CLDN4 marks the T2 substate in AT2-less in situ regions and KRT8^high^ marks T1 through T3 substates. KRT8^high^ cells in de novo regions lack CLDN4. (**B**) KRT8^high^ cells in de novo regions are T1 (HOPX-) or T4 (HOPX+KRT8-). (Scale bar 50µm).

**Fig. S3.**
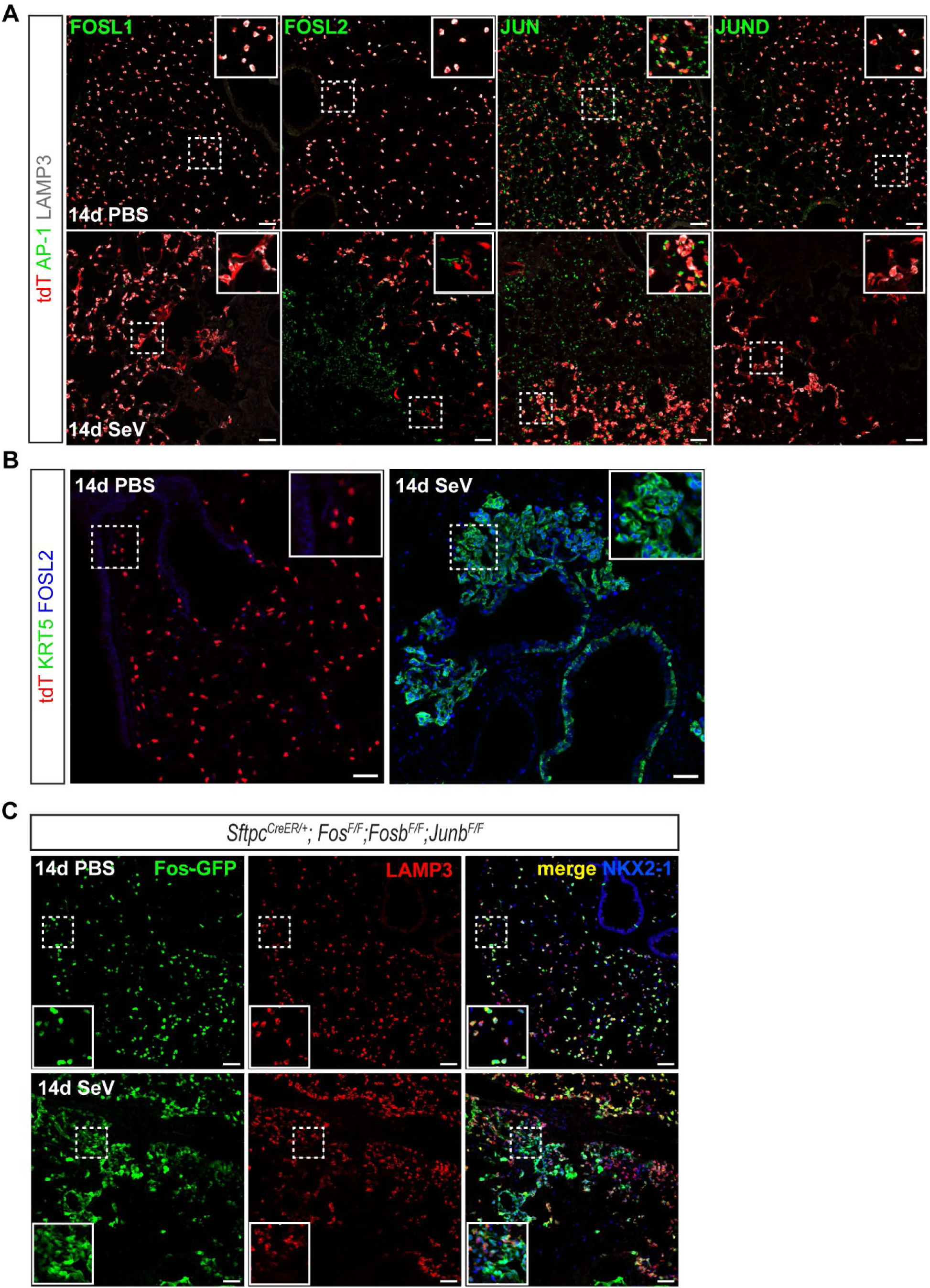
Expression of AP-1 members FOS, FOSL1, FOSL2, JUN, and JUND. (**A-B**) Confocal images of 10µm sectioned lungs from PBS or SeV infected *Sftpc^CreER/+^;Rosa^tdT/+^* mice, showing either absence (FOSL1, FOSL2, JUND) or constant (JUN) expression in lineage-traced AT2 cells (tdT+) upon infection (**A**). FOSL2 is present in airway cells at baseline and airway-derived KRT5+ pods upon infection (**B**). (**C**) Confocal images of 10µm sectioned lungs from PBS or SeV infected *Sftpc^CreER/+^*;Fos*^F/F^*;Fosb*^F/F^*;Junb*^F/F^*mice. Fos-GFP is expressed upon *Sftpc^CreER^* recombination and under the control of its endogenous promoter/enhancer, serving as a surrogate of FOS antibody staining as no reliable FOS antibody works in the lung. Fos-GFP is in all AT2 cells (LAMP3+) at baseline (PBS) as well as transitioning AT2 cells (LAMP3-NKX2-1+) upon infection. (Scale bar 50µm).

**Fig. S4.**
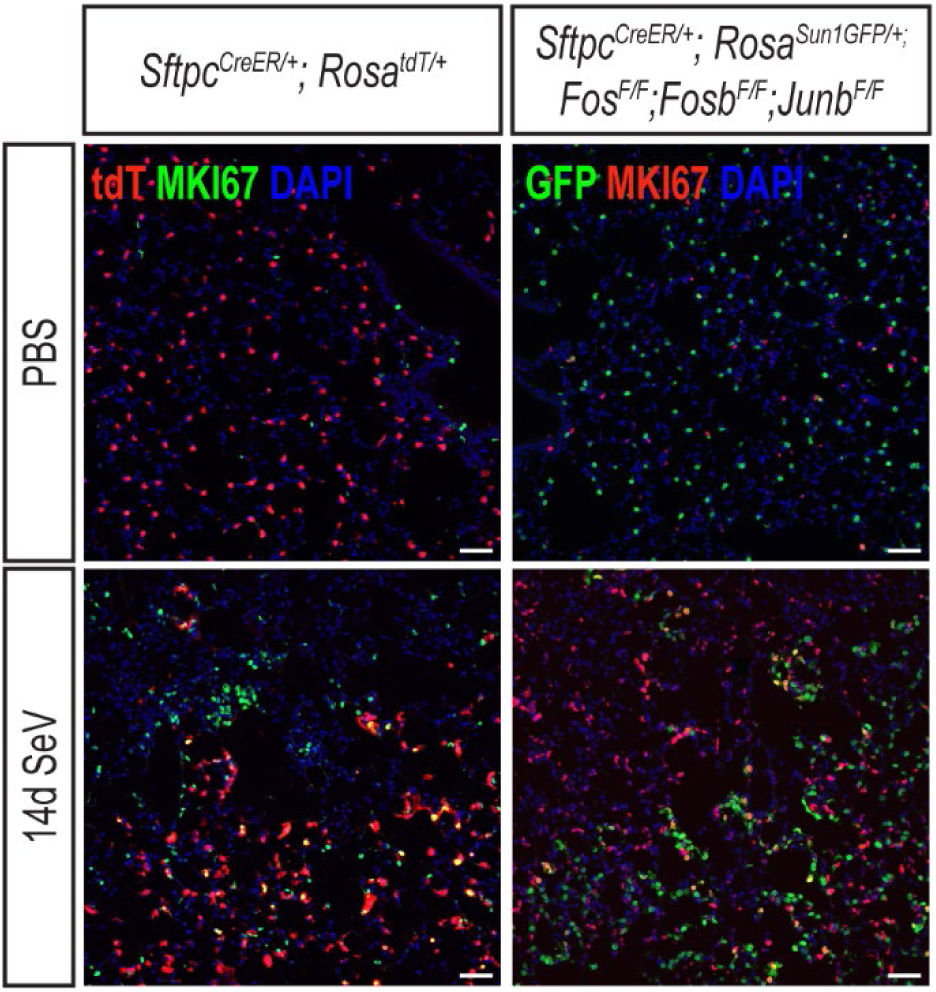
Comparable proliferation in AT2 cells of infected wildtype and AP-1 mutant lungs. Confocal images of 10µm sectioned lungs from PBS or infected *Sftpc^CreER/+^;Rosa^tdT/+^*and AP-1 mutant mice immunostained for a proliferation marker MKI67. (Scale bar 50µm).

**Fig. S5.**
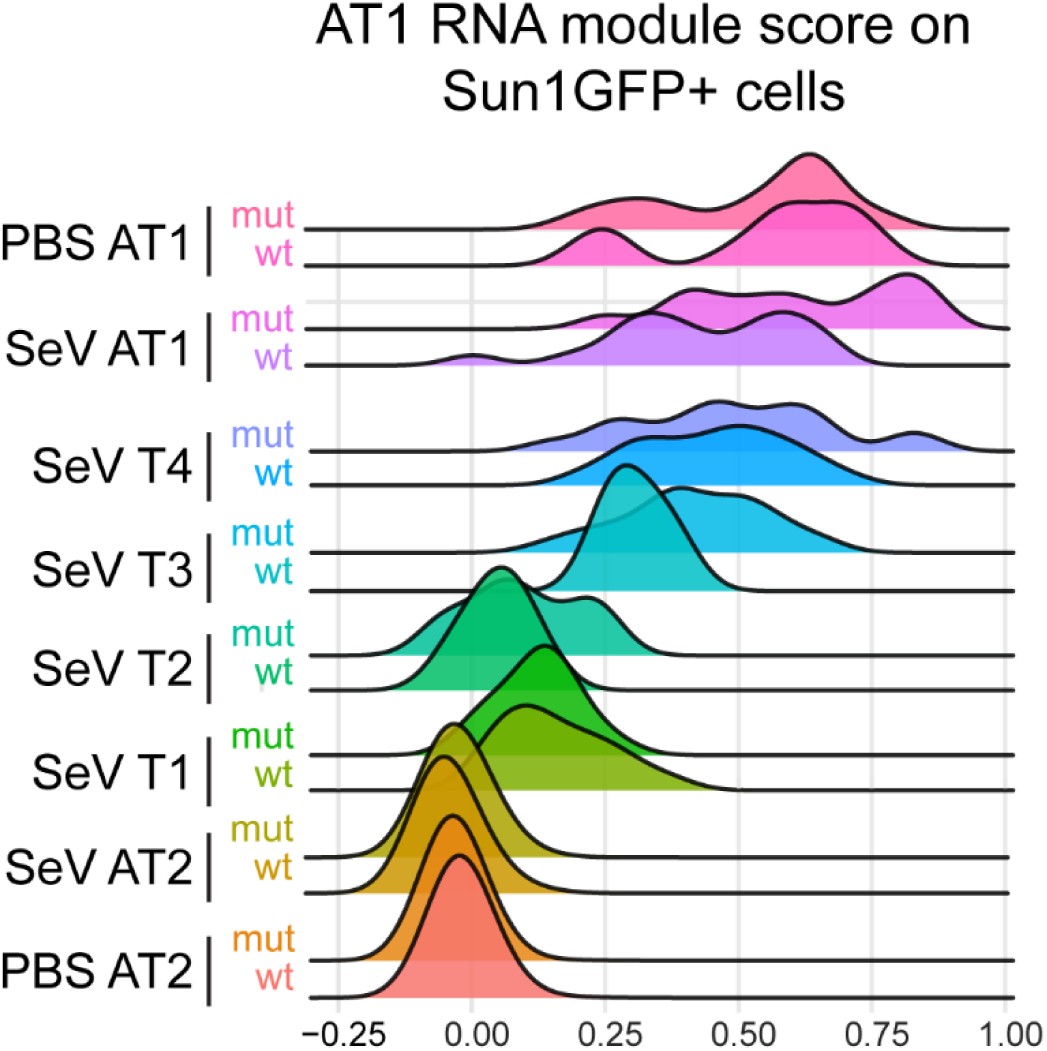
AP-1 mutant populations have enhanced overall AT1 differentiation. Single-cell multiome ridgeplot of AT1 RNA module score for AT2-derived SunGFP^+^ cells to remove the confounding effect of baseline AT1 cells.

